# A pilot study on the home range and movement patterns of the Andean Fox *Lycalopex culpaeus* (Molina, 1782) in Cotopaxi National Park, Ecuador

**DOI:** 10.1101/2021.02.10.430628

**Authors:** Armando Castellanos, Francisco X. Castellanos, Roland Kays, Jorge Brito

## Abstract

This study reports movement patterns and home range estimates of an Andean fox (*Lycalopex culpaeus*) in Cotopaxi National Park in Ecuador, representing the first GPS-tagging of the species. The GPS functioned well during the 197-day tracking period. Home range sizes ranged between 4.9 - 8.1 km^2^, depending on the estimation method. Movement speeds averaged 0.17 km/hr at day vs. 0.87 km/hr. at night, and distance traveled averaged 0.23 km at day vs. 0.89 km at night. These preliminary results highlight the importance of collecting unbiased, high-quality data which enables an enhanced understanding on mammal behavior and human/animal interaction.

---

The Andean Fox *Lycalopex culpaeus* Molina 1782 is native to western South America, distributed from southern Colombia to southern Argentina (Ramírez-Chaves et al. 2013). It is the second-largest canid in South America (Sillero-Zubiri 2009), with nocturnal and crepuscular habits (Monteverde and Piudo 2011). The Andean fox is an opportunist that typically eats small mammals but can also feed on fruits (Cadena-Ortíz et al. 2020).

Studies of the home range of the Andean Fox have found sizes of 3.8–18.8 km^2^ in females and 1.4–7.3 km^2^ in males (Table 1). However, such studies have been hitherto absent in Ecuador and only VHF tracking collars have been used in a small part of its wide distribution range. Herein, we report the first use of a GPS collar for this species and movement analysis of an Andean Fox in Ecuador.

**Table 1.**
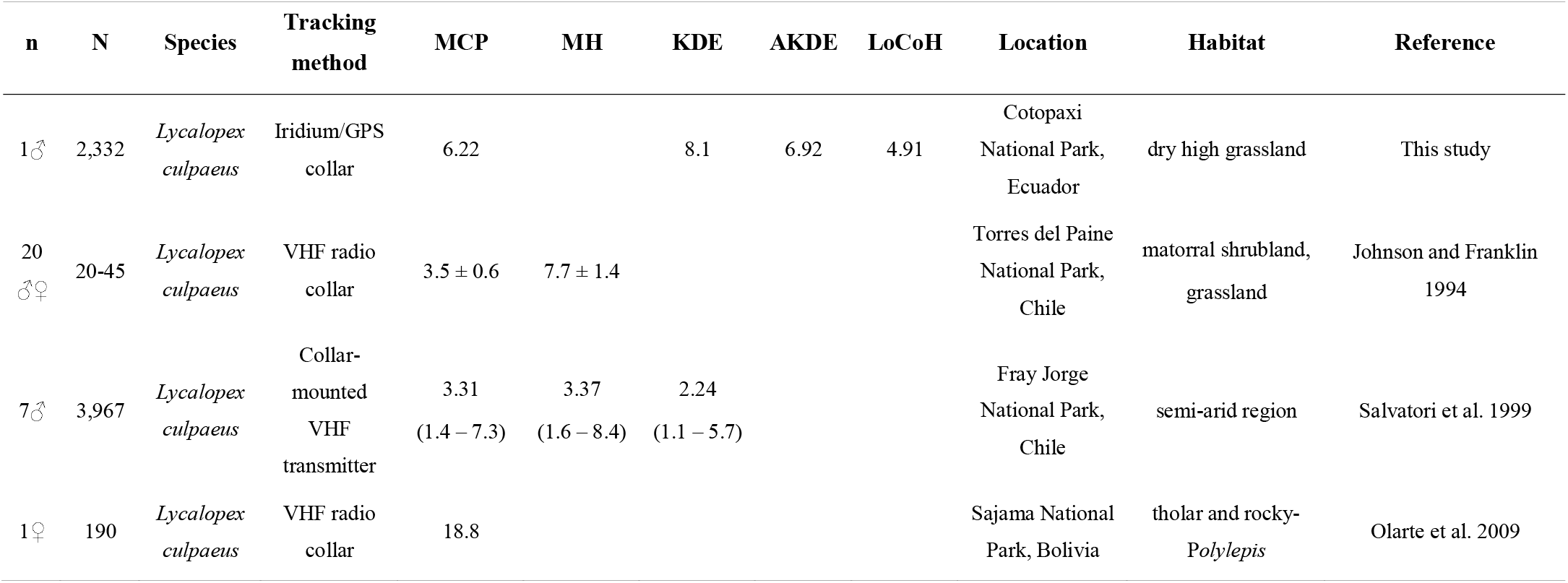
Male and female seasonal home range sizes measured in km^2^. To facilitate comparison with our 6-month tracking period analysis, only areas enclosing the 95% of habitat utilization are shown, except for Salvatori et al. (1999) 100% MCP. n = sample of adult males or females; N = recorded locations; MCP = Minimum Convex Polygon; MH = Mean harmonic; KDE = Kernel Density Estimate; AKDE = Autocorrelated Kernel Density Estimate; LoCoH = Local Convex Hull.

| n | N | Species | Tracking method | MCP | MH | KDE | AKDE | LoCoH | Location | Habitat | Reference |
| --- | --- | --- | --- | --- | --- | --- | --- | --- | --- | --- | --- |
| 1♂ | 2,332 | *Lycalopex culpaeus* | Iridium/GPS collar | 6.22 |  | 8.1 | 6.92 | 4.91 | Cotopaxi National Park, Ecuador | dry high grassland | This study |
| 20 ♂♀ | 20-45 | *Lycalopex culpaeus* | VHF radio collar | 3.5 ± 0.6 | 7.7 ± 1.4 |  |  |  | Torres del Paine National Park, Chile | matorral shrubland, grassland | Johnson and Franklin 1994 |
| 7♂ | 3,967 | *Lycalopex culpaeus* | Collar-mounted VHF transmitter | 3.31<br>(1.4 – 7.3) | 3.37<br>(1.6 – 8.4) | 2.24<br>(1.1 – 5.7) |  |  | Fray Jorge National Park, Chile | semi-arid region | Salvatori et al. 1999 |
| 1♀ | 190 | *Lycalopex culpaeus* | VHF radio collar | 18.8 |  |  |  |  | Sajama National Park, Bolivia | tholar and rocky-*Polylepis* | Olarte et al. 2009 |

On 26 September 2019 we captured an adult male Andean Fox in the Cotopaxi National Park (0° 33′ 42.1″ S 78° 26′ 00.5″ W; 3,647 m.a.s.l; Fig. 1A) with a wire box trap. The animal was immobilized with an intramuscular dart with a combination of Ketamine hydrochloride and Xylazine hydrochloride (Castellanos et al. 2020), weighed (10.4 kg), measured (head length = 21 cm.; body length = 65 cm.; neck circumference = 29 cm.; chest circumference = 41 cm.; and tail length = 53 cm.) and assessed its health status (healthy). This specimen was fit with an Iridium/GPS collar, Lite model of 450 g., (Vectronic Aerospace GmbH, Berlin, Germany), which also was equipped with a VHF transmitter, temperature sensor, mortality sensor, and a drop-off mechanism. Capture was performed through scientific research authorization No MAAE-ARSFC-2020-0642, and guidelines of the Ministry of Environment of Ecuador for the use and care of animals were followed.

**Figure 1.**
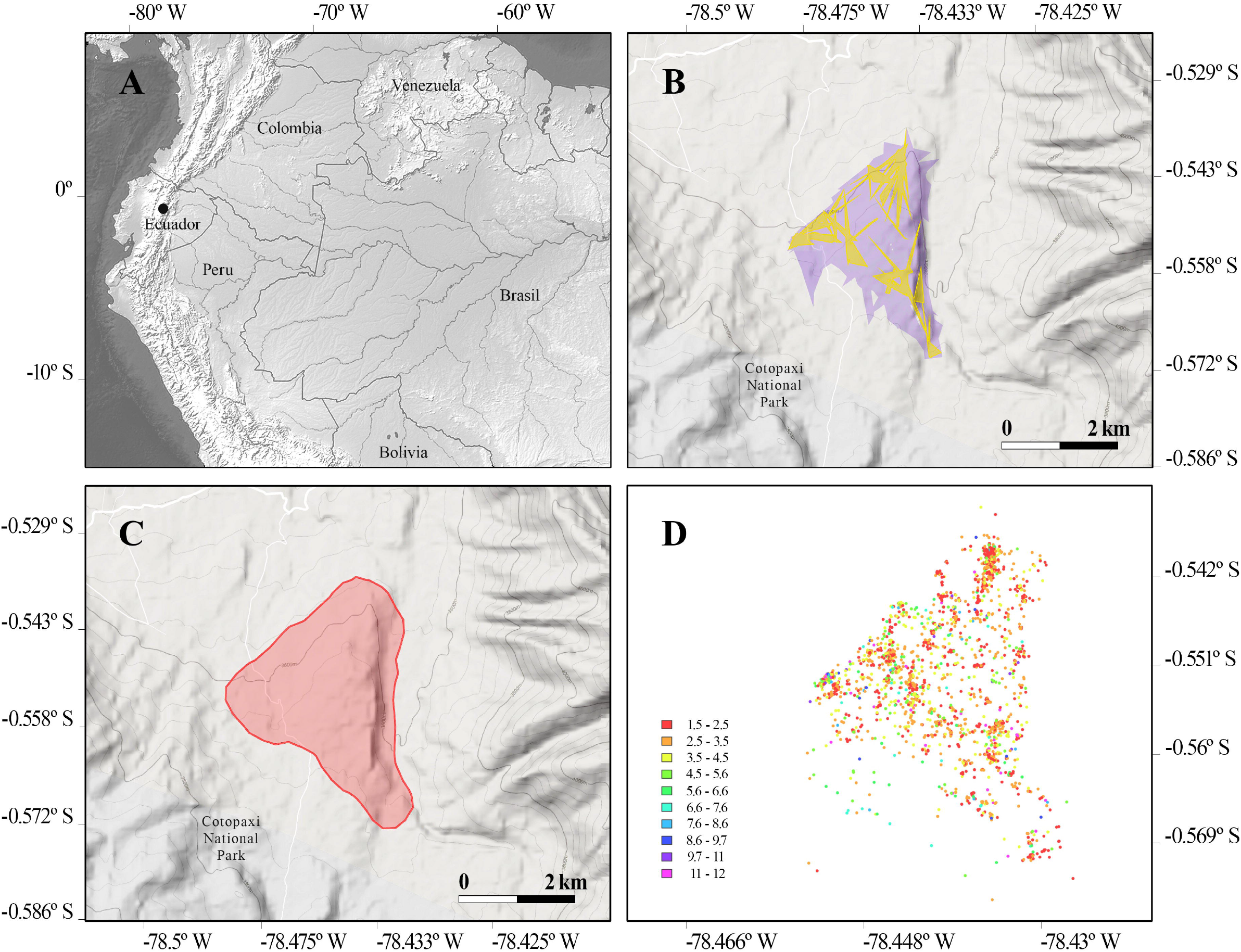
A) Study site located in Cotopaxi National Park B) Home range (violet) and core area (yellow) of Mashca created from 2,332 locations using the fixed k-method. C) 95% KDE D) Mashca’s fixed positions colored according to the mean number of locations per visit. Color time representations are shown to the left and measured in hours. Hulls were created from 2,329 locations using the fixed k-method.

This fox, nicknamed “Mashca” was monitored from September 26, 2019 to April 10, 2020 (197 days) until its death, due to unknown reasons. GPS sampling rates were set to two hours. An immediate retrieval of this specimen’s collar after death was not possible due to transportation restrictions in effect due to the COVID-19 pandemic in Ecuador. However, these stationary GPS positions taken following Mashca’s death were used as opportunistic calibration data to calculate the location error for the collar (Fleming et al. 2020).

GPS fixes were successful in 2,332 of 2,367 attempts (98.5%). To filter out low quality fixes 35 GPS positions with DOP > 3 values and 2D-3D GPS fix types were removed, leaving 2,332 high quality locations. Moreover, three positions were manually marked as outliers since they were evident incorrect coordinates based on impossibly fast movements. The dataset analyzed in this study is available in the Movebank Data Repository https://doi.org/10.5441/001/1.1q2c075p (Castellanos 2021).

Home ranges were estimated using four different methods: Minimum Convex Polygon (MCP), Kernel Density Estimate (KDE), Autocorrelated Kernel Density Estimate (AKDE), and Nearest-Neighbor Convex Hull k-method (k-NNCH) (Fleming and Calabrese 2017; Getz and Wilmers 2004; Lyons et al. 2013; Mohr 1947; Seaman and Powel 1996) using R v4.0.0 (R Core Team 2020), and the adehabitatHR, and T-LoCoH packages (Calabrese et al. 2016; Calenge 2006; Lyons et al. 2018). The latter was also used to examine hull metrics for time-use dimensions in order to identify habitat usage and behavior patterns. An inter-visit gap period of ≥12 hours was defined to obtain the average duration of each visit (mnlv; mean number of locations per visit), and visitation rates to the hulls (nsv; number of separate visits).

The 95% MCP method resulted in a 6.22 km^2^ home range. The 95% KDE estimate was 8.1 km^2^ (Fig. 1C) while the 95% AKDE estimates was 6.92 km^2^. Lastly, the T-LoCoH package allowed to estimate a 95% home range of 4.91 km^2^ (Fig. 1B). We also estimated the core area using the 50% AKDE (1.99km^2^) and 50% T-LoCoH isopleths (1.53km^2^). These core areas overlapped with sites of high anthropogenic activity, especially in Los Mortiños Hostal and agricultural fields. In general, the fox moved around its home range without staying for more than ~5 hours in one area (Fig. 1D). Duration of visit (mnlv) around the crops was of ~1–5 hours and visitation rates (nsv) estimate that hulls were visited from ~60–140 times. Likewise, mnlv around Los Mortiños Hostal indicate that visits didn’t last longer than ~7 hours, although Mashca stayed over 10 hours few times, and nsv were lower than around crop areas mostly between ~40–80 times (Fig. 1D).

Our data supports the current knowledge of the nocturnal and crepuscular activity of this species. Distances and speed were greater during night hours, probably due to increasing foraging and hunting behaviors (Fig. 2). In daylight, Mashca traveled minimum and maximum distances of 0 and 2.7 km, respectively (average 0.23 ± 0.39 km), whereas at night it traveled minimum distances of 0 km and maximum of 3.34 km (average 0.89 ± 0.7 km). Average speed during the whole tracking period in daylight was 4.08 km/day, while instantaneous speeds estimates were 0.17 ± 0.06 km/hr. with minimum speeds around 0.14 km/hr. and maximum of 0.63 km/hr. At night, the average speed of the complete sampling period was 20.8 km/night, and instantaneous speeds were around 0.87 ± 0.09 km/hr. with minimum and maximum speeds of 0.76 – 1.35 km/hr. respectively (Fig. 2).

**Figure 2.**
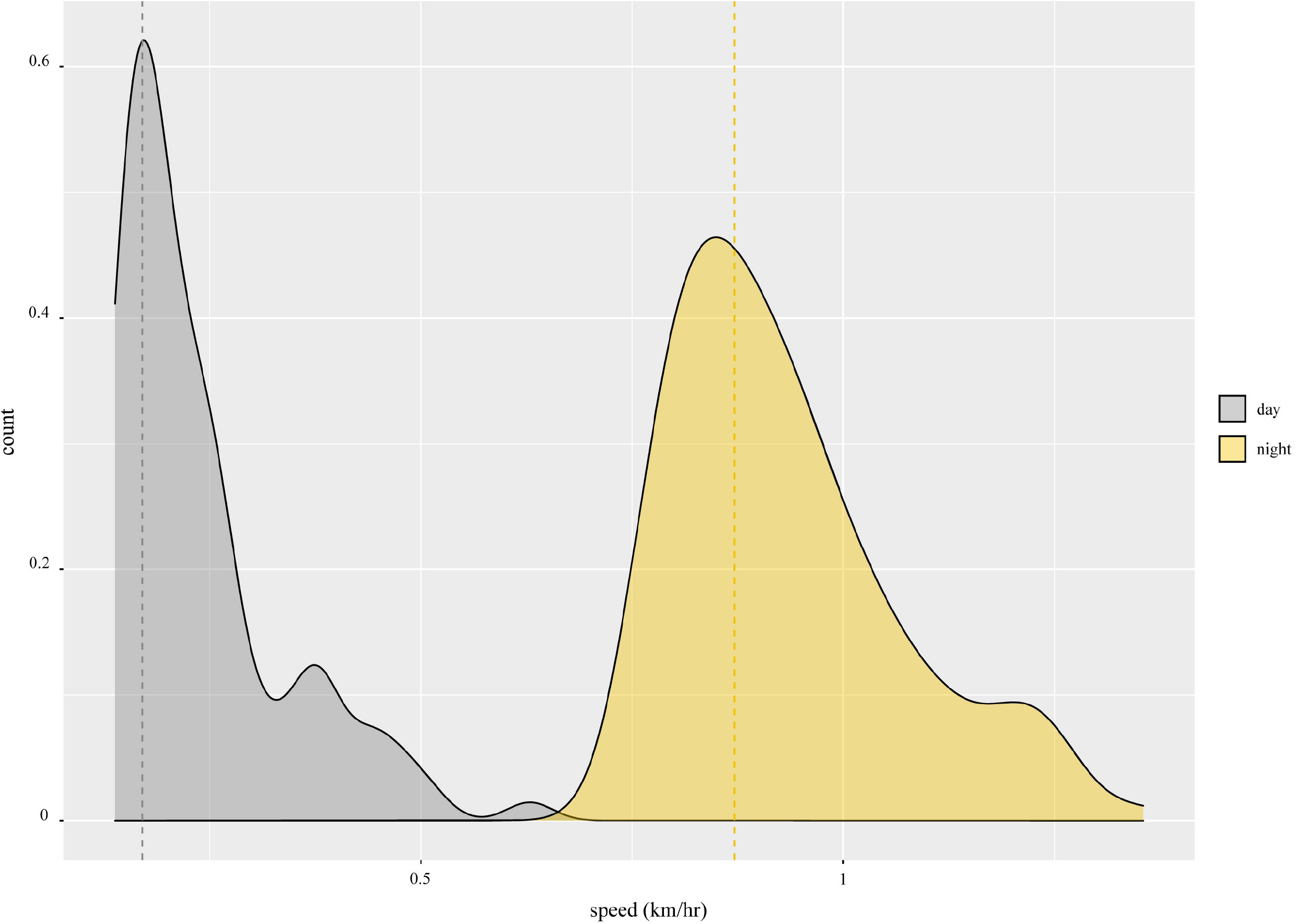
Instantaneous speeds estimates obtained through the ctmm package. Mashca’s speeds during the day and night are shown in gray and yellow, respectively. Dotted lines show average speeds.

This is the first GPS collaring of this species, showing that these new small tracking units can collect useful data in dry mountainous terrain, with an accuracy of 35 m. A combination of older (MCP) and newer (AKDE) home range methods were used to facilitate comparison with previous studies. Our estimates for home range size for the Ecuadorian Andean fox are larger than the ones obtained in other studies (Table 1). We suspect some of this difference may be due to the difficulty of recording VHF locations compared to GPS, even in highly similar habitats (Table 1). Indeed, Olarte et al. (2009) had 190 fixes in a single female, while Johnson and Franklin (1994) had a minimum of 20 and a mean of 45 observations per individual (Table 1). In comparison, we obtained 2,332 high quality locations in 6 months, with very few missed fixes, obtaining an unbiased and comprehensive measure of animal movement, compared to VHF tracking.

The AKDE and T-LoCoH core and home range areas presented in this study are the first for this species. Interestingly, core areas overlap highly anthropized areas (Fig. 1B). On one hand this suggests that the species is adapted to human disturbance. However, the early mortality of this individual might also show the riskiness of this strategy (although cause of death could not be determined).

Even though no strong conclusions can be drawn from the data analysis of a single male specimen, it is noticeable that home ranges estimated in our study are different from several specimens of different South American countries, probably due to the high number and high quality of locations recorded (Table 1). Similar results have also been found when comparing MCP and KDE methods with VHF vs GPS technologies in other animals, such as omnivores (*Ursus arctos*; Arthur and Schwartz 1999), carnivores (*Panthera tigris altaica*; Hernandez-Blanco et al. 2015), and herbivores (*Odocoileus virginatus*; Kochanny et al. 2009). Furthermore, the unbiased nature of our data allows to undoubtedly conclude on the nocturnal behavior of this specimen.

With the success of this pilot study, additional work should continue to expand the knowledge of this species across its range, incorporating satellite tracking technology and modern analytical methods to estimate animal movement and home ranges in order to learn more about their interrelationships with other species, and with humans.

## Acknowledgements

We thank the Cotopaxi Provincial Office/MAAE, Cotopaxi National Park and their park rangers, Watershed and Water Protection Fund (FONAG), Wilson Lutuala for the help provided, Susana Escandón for the research sponsorship, and Angel Yánez-Zapata for his assistance in field work. Special thanks to Christen Fleming and Shauhin Alavi for their help with the ctmm package.

## Author contributions

Armando Castellanos and Jorge Brito conceived this study and gathered data. Francisco Castellanos analyzed data guided by Roland Kays. All authors wrote and reviewed the manuscript prior to submission.

## Research funding

The procurement of the collars has been funded by the International Climate Initiative (ICI or IKI in German) of the German Federal Ministry for the Environment, Nature Conservation, Building and Nuclear Safety with support of KfW Development Bank (BMZ project no. 2098 10 987).

## Compliance with ethical standards

Handling and all activities regarding specimens followed ethical procedures recommended by the American Society of Mammalogists (Sikes et al. 2016).

